# SegBio: A lightweight end-to-end toolkit for Instance Segmentation of biological samples

**DOI:** 10.64898/2026.04.03.716031

**Authors:** Eduard Bokman, Neta Barlam, Omri Babay, Yuval Balshayi, Yifat Eliezer, Alon Zaslaver

## Abstract

High-throughput phenotyping of biological samples is essential for large-scale studies but is frequently bottlenecked by the need for accurate instance segmentation in crowded images. While deep learning offers powerful solutions, the high cost of manual annotation and the requirement for coding expertise often limit adoption in routine laboratory workflows. Here we present SegBio, a lightweight, open-source pipeline that enables end-to-end instance segmentation for non-expert users. The protocol features an interactive annotation GUI that extrapolates full masks from minimal centerline markings, significantly reducing manual labeling effort. It further integrates a configurable U-Net training module and a standalone inference application with a ‘human-in-the-loop’ editing workflow for rapid and intuitive error correction. We employ the pipeline to annotate and train the model on a novel dataset of crowded *C. elegans* images. Validated on independent datasets, SegBio achieves high segmentation performance (Panoptic quality ∼0.85) and accurately quantifies per-animal morphology and fluorescence. By eliminating external dependencies and streamlining the correction process, SegBio provides a scalable solution for routine phenotyping that is easily generalized to other crowded biological samples, such as cellular organelles, cells, and organisms.

## Introduction

Deep learning (DL) approaches, particularly U-Net–style architectures and their variants, have become a standard choice for biomedical semantic segmentation because they are relatively efficient and can generalize well when trained on sufficiently diverse annotated data ^1–3^. At the same time, crowded microscopy often requires instance segmentation, and a broad landscape of solutions has emerged, including detection-first approaches ^4–6^, shape-representation methods tailored to microscopy ^7–9^, and generalist “off-the-shelf” tools designed for broad transfer across imaging conditions ^10,11^. Many practical pipelines also combine CNN probability maps with classical post-processing (such as marker-controlled watershed) to split touching objects into individual instances ^12–14^. Nevertheless, several practical obstacles continue to limit adoption in many labs.

Training a DL model from scratch requires generating large annotated datasets, which can be prohibitive if done manually. On the other hand, pre-trained models and public datasets are not always usable off-the-shelf due to differences in image setups between labs (microscopes, acquisition settings, sample preparation, etc.). Even modest distribution shifts can degrade performance and force users to fine-tune or retrain models on locally acquired images ^10,15^. Additionally, even strong models are not fully autonomous in practice. Noisy samples, and occasional merge/split errors and other edge-case failures still require human supervision and targeted manual edits to maintain a high standard of accuracy, making usability and correction workflows crucial for lowering the barrier for non-expert users while preserving reproducibility and transparency ^16,17^.

*Caenorhabditis elegans* is a widely used model organism for studying development, aging, disease mechanisms, and gene function, in part because it is small, optically accessible, and amenable to scalable imaging assays ^18–22^. Large-scale phenotyping in brightfield and fluorescence microscopy often depends on extracting per-animal measurements (e.g., count, size, posture-derived metrics, or fluorescence intensity) rather than relying on whole-image summary statistics ^23–26^. However, obtaining these measurements typically requires accurate segmentation of individual worms, a step that can become a major bottleneck when experiments generate a large number of images, or when animals are tightly clustered.

*C. elegans* computer vision has long included behavioral-analysis pipelines that do not rely on dense instance masks, ranging from early machine-vision tracking and phenotype classification to newer segmentation-free methods for locomotor quantification under noisy imaging conditions ^22,23^. However, manual instance segmentation remains common in worm imaging pipelines because it provides high-quality masks and can be used directly for downstream morphology and fluorescence quantification ^27,28^. Yet manual outlining is slow, labor-intensive, and introduces variability across users and labs. This challenge is amplified in crowded microscopy fields where adjacent worms are difficult to separate consistently. There is, therefore, a strong incentive to develop automated or semi-automated worm segmentation tools, enabling high-throughput image analysis, while maintaining accuracy through human-in-the-loop editing workflows.

Related challenges also arise in whole-brain *C. elegans* imaging, where automated segmentation, correspondence, and tracking of densely packed neurons in moving animals have motivated pipelines based on registration, semi-synthetic training, targeted augmentation, and multi-lab data harmonization ^29–32^.

Here we introduce SegBio, an open-source, lightweight instance segmentation tool, pre-trained for crowded brightfield microscopy images of adult *C. elegans*, and designed to work as a simple, intuitive off-the-shelf solution for routine use. At the same time, SegBio includes three integrated modules that directly address the practical obstacles above: (i) a fast annotation module to reduce the cost of generating lab-specific training data, (ii) a training module that supports straightforward retraining or fine-tuning the model to novel imaging conditions, and (iii) a standalone inference-and-editing workflow that enables efficient human-in-the-loop correction of model outputs.

We first provide an overview of the pipeline, followed by a detailed implementation and usage guide for each component. We then provide quantitative validation of the pre-trained model on two independently curated validation sets to estimate out-of-sample performance and typical correction burden. Finally, we demonstrate practical common downstream analyses enabled by the resulting instance masks, including per-animal morphology measurements and fluorescence quantification.

In summary, SegBio provides an accessible and flexible image analysis pipeline that lowers the barrier to adopting deep-learning segmentation in everyday experiments, supporting reproducible, scalable worm phenotyping from raw microscopy images to analysis-ready measurements.

## Results

### Software overview

SegBio provides a simple, open-source end-to-end solution for instance segmentation, from training data creation by manual annotation to deep-learning based automatic labeling and morphological feature extraction. The toolset presented here is fine-tuned for adult *C. elegans* nematodes but is easily adaptable for a wide range of other use-cases.

The library includes three software modules, including two simple and intuitive Graphical User Interface modules (GUIs):

i. Manual annotation GUI for fast labeling/data curation (Fig 1A)
ii. CNN-based (U-Net) instance segmentation training library (Fig 1B)
iii. Automatic inference module and manual editing GUI (Fig 1C)

**Figure 1.**
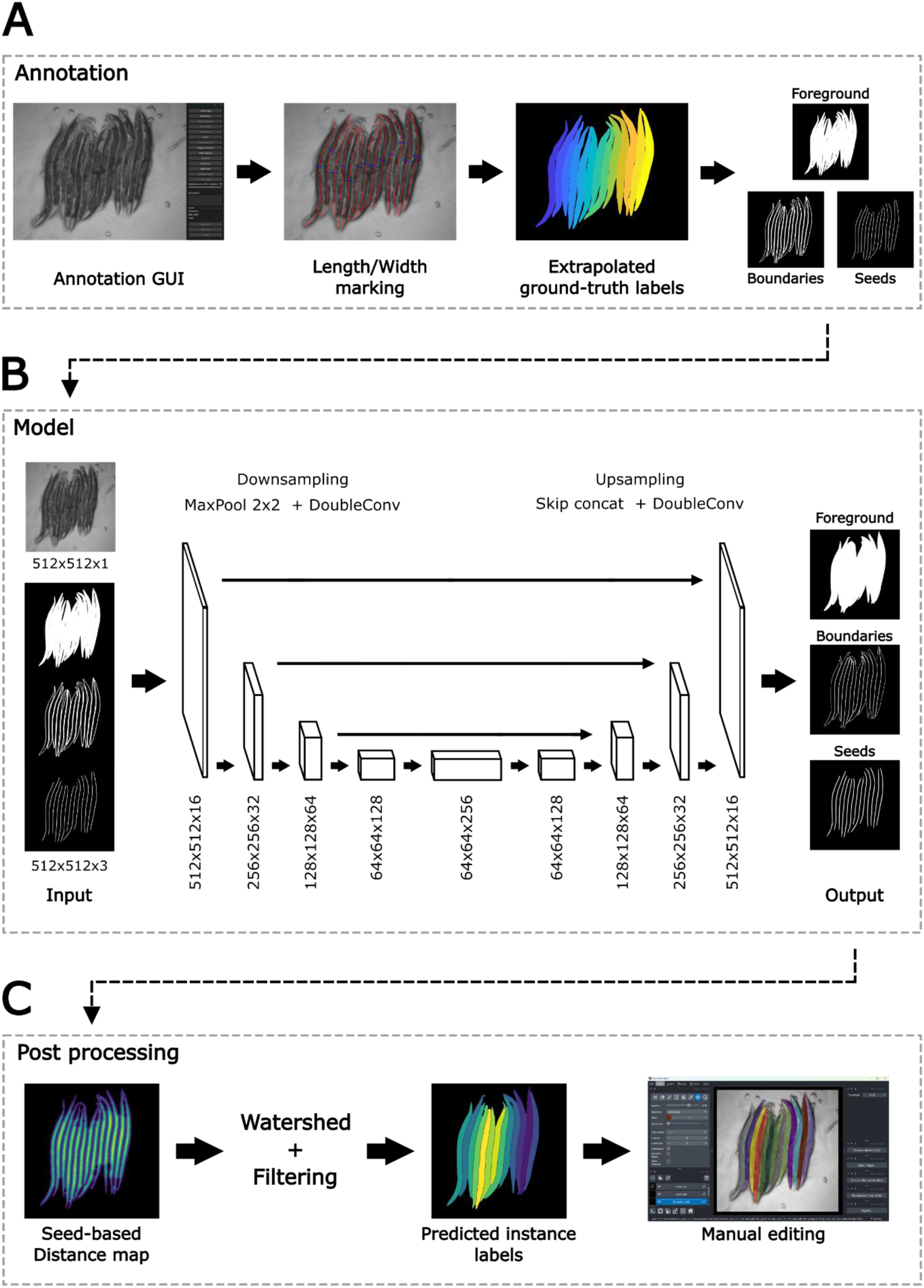
The segmentation pipeline consists of annotation, model training, and inference modules. **A)** The annotation module (i) is an interactive manual marking interface where the user marks the length and width of the object. Full instance masks are then extrapolated from the markings. **B)** The model training module (ii) consists of a configurable U-Net that performs a 3-class semantic segmentation on all object pixels (foreground), object outlines (boundaries), and object centerline (seeds). The separate class masks are obtained by deconstruction of the object masks from module 1. The U-Net is constructed with flexible depth and width, allowing users to easily adapt and retrain the model for their own data. **C)** The inference module (iii) provides a simple and intuitive GUI to perform automatic instance segmentation of images. The 3-class predictions given by the model are transformed into instance masks via a watershed algorithm that fills in objects growing outward from the seeds until a boundary is reached. The resulting object masks are then displayed in the GUI on top of the image for simple manual editing. The user can apply quick fixes by lightly adjusting model outputs and rerunning the post-processing segmentation. Alternatively, the user can edit the masks themselves for finer control.

The first two modules provide open-source data curation and model training pipelines with which the toolset can be adapted for additional tasks. Module (i) is a standalone, self-contained, Python-based executable GUI that allows users to create full instance masks of elongated objects without painstakingly tracing the entire outline of the object. The user must only trace the centerline, specify the widest point of each object, and supply a characteristic tapering profile. The software then extrapolates this data into full object masks. This method is much faster and requires significantly less tracing accuracy than hand-drawing a full outline.

Module (ii) is a PyTorch-based flexible U-Net training pipeline that can be used to retrain the model on new data. Users can easily adjust multiple parameters to better fit the model for their needs, including the architecture of the network (number of layers and filters), image augmentations, post-processing method and filtering criteria. The resulting model can then be integrated into module (iii) for inference.

The inference GUI (iii) is delivered as a standalone, self-contained executable with pre-trained model weights tuned for crowded microscopy images of adult *C. elegans*. The package is written in Python (PyTorch), but does not require an existing Python installation, has no external dependencies, and can employ either GPU or CPU for inference. As such, the GUI is highly accessible to anyone, regardless of available hardware or coding knowledge, and can be used out-of-the-box for multiple applications in *C. elegans* research, including counting individuals, extracting morphological measurements, and quantifying fluorescence.

Detailed user and developer guides for all modules are available in https://github.com/zaslab/SegBio.

### Model validation

To assess the quality of the model’s predictions we used two validation sets created separately from the training set by two different experimenters. The experimenters used the model to segment images and then manually adjusted the output with the editing tool. We then compared the raw predictions of the model to the final user-adjusted masks (Fig 2, Table 1). This provides an estimate of the independent accuracy of the model on new and variable data, and the amount of required user post-processing.

**Figure 2.**
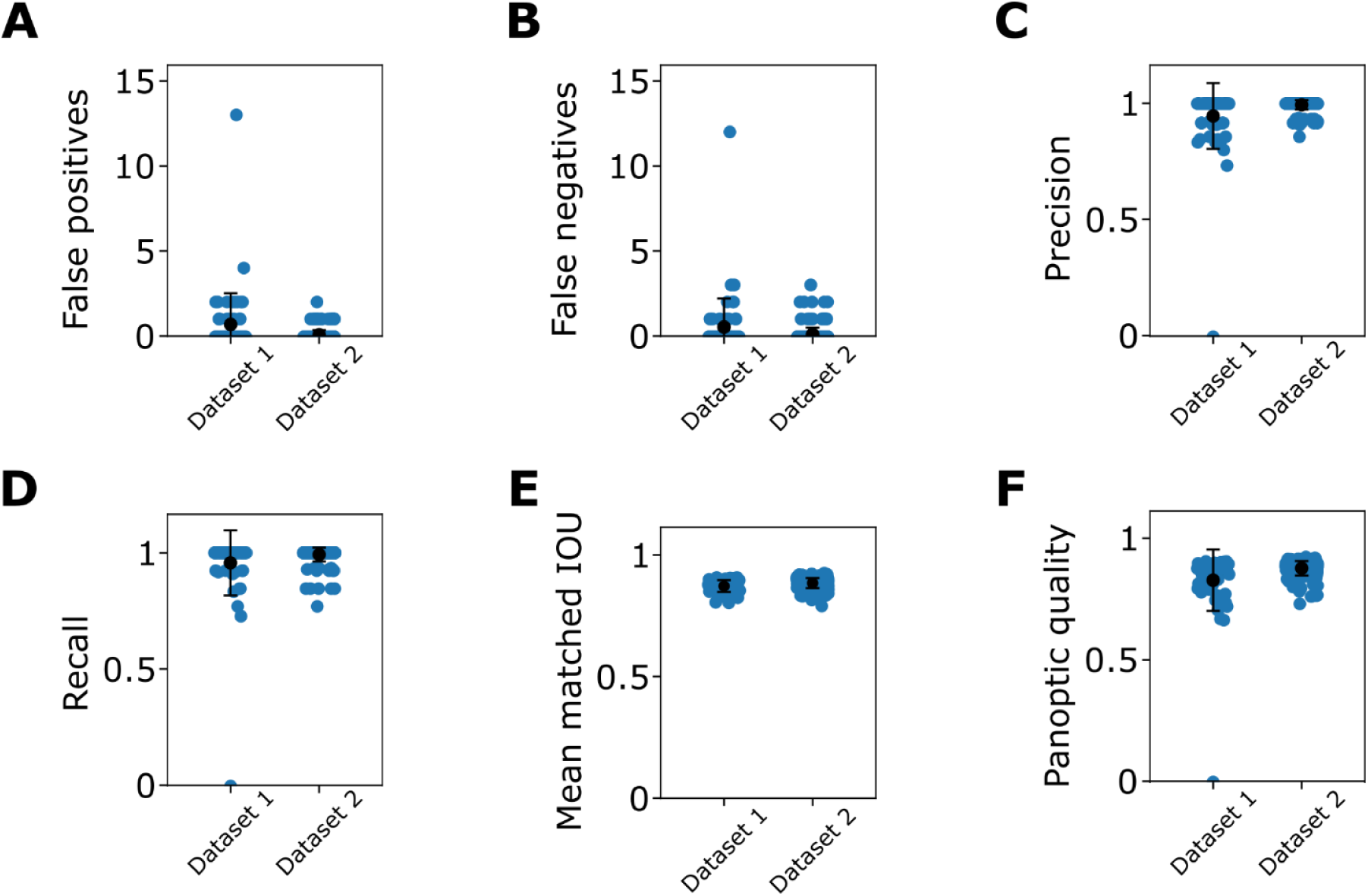
The model accurately segments individual worms with minimal manual editing. **A)** False positive per image, matched at a threshold of IoU>0.5. False positives normally occur when a worm is over-segmented into multiple individuals. Such fragmentations can be fixed by removing the extra boundaries separating the fragments. **B)** False negatives per image, matched at a threshold of IoU>0.5. False negatives can occur when a full worm is either not segmented or is filtered out during post processing. These can be mitigated by adjusting the filtering criteria or by manually adding a new mask. **C)** Precision per image. **D)** Recall per image. **E)** Intersection over union for individual worm masks, averaged over each image. **F)** Panoptic quality per image, a summary statistic combining object detection and mask overlap (recognition quality and segmentation quality, respectively, Table 1).

**Table 1.** Model performance metrics on two separate test sets.

| Dataset | Dataset 1 |  | Dataset 2 |  |
| --- | --- | --- | --- | --- |
| N images | 60 |  | 250 |  |
| Metric | Mean | Std | Mean | Std |
| N ground truth | 12.339 | 1.108 | 13.558 | 0.971 |
| N predicted | 12.492 | 1.104 | 13.530 | 1.021 |
| Objects found (%) | 0.957 | 0.140 | 0.993 | 0.030 |
| True positive | 11.814 | 2.047 | 13.458 | 1.051 |
| False positive | 0.678 | 1.842 | 0.072 | 0.274 |
| False negative | 0.525 | 1.685 | 0.100 | 0.392 |
| Precision | 0.947 | 0.142 | 0.995 | 0.020 |
| Recall | 0.957 | 0.140 | 0.993 | 0.030 |
| Recognition quality | 0.951 | 0.138 | 0.993 | 0.022 |
| Segmentation quality | 0.872 | 0.024 | 0.885 | 0.020 |
| Panoptic quality | 0.830 | 0.126 | 0.879 | 0.030 |
| Matched instance IoU | 0.872 | 0.024 | 0.885 | 0.020 |
| Foreground IoU | 0.866 | 0.065 | 0.891 | 0.022 |

Across the two test sets, the model performed consistently well in both object detection, and mask matching. From the perspective of object detection, the two main failure modes of the model are over-segmentation, where a single object is divided into multiple instances (false positives, fp), and under-segmentation, where multiple objects are combined into a single instance (false negatives, fn). The model performed well on both metrics, correctly identifying the vast majority of worms in an image (Fig 2 A-D). The number of both over- and under-segmented instances (fp and fn, respectively, Fig 2 A,B) in both test datasets was less than one per image, resulting in precision and recall of ∼1 (Fig 2 C,D). For correctly identified objects, quality of segmentation was also high, with an average IoU of ∼0.88 (Fig 2 E) across datasets. This indicates strong overlap between predicted masks and ground truth. Panoptic quality, a summary metric combining object detection and mask overlap, was ∼0.85 (Fig 2 F) across datasets, indicating strong overall performance of the model.

### The toolset supports multiple applications of worm segmentation

Many research applications require accurate morphological measurements of individual animals. These include studies on development, aging, diseases, etc. Manual tracing of individuals is slow, labor intensive, and can suffer from experimenter bias. Our model solves these issues by quickly and accurately creating masks of single animals, and extracting common and useful metrics from these masks including object length, width, and area, as well as allowing for quantification of fluorescence intensity in transgenic animals. Further measurements (*e.g*. posture, tapering) can be obtained from the masks as needed.

Figure 3A shows microscopy images obtained with an additional fluorescence channel of *hsp-6::GFP* worms that exhibited elevated fluorescence after consuming bacteria exposed to zinc in a concentration dependent manner (see methods). The images were segmented using the trained model and inference GUI, and the predicted masks were used to quantify the median fluorescence intensity of all the pixels within each worm. The masks provide an accurate measurement of fluorescence, capturing the dose-dependent GFP response (Fig 3B).

**Figure 3.**
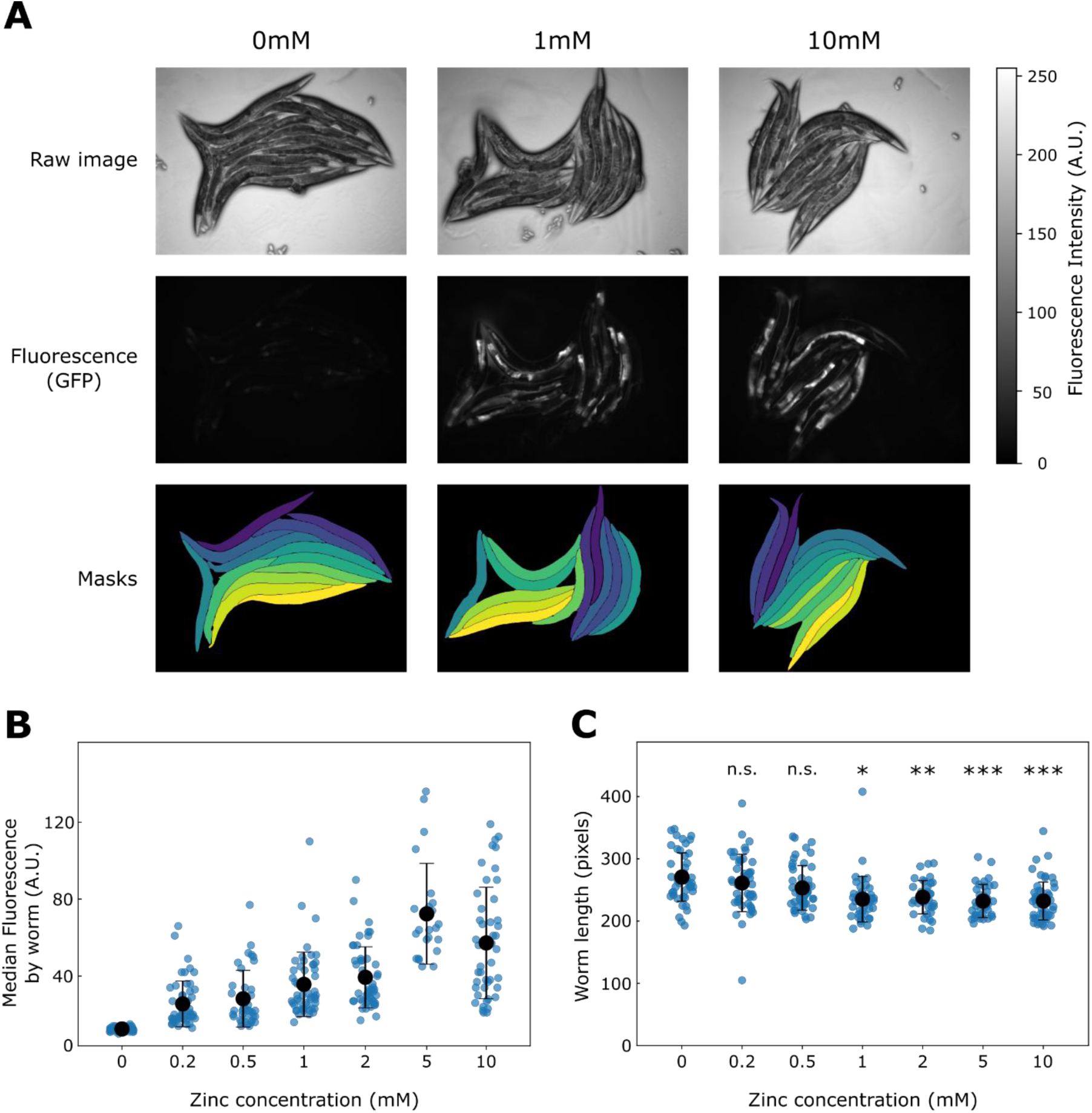
SegBio can be employed to measure fluorescence intensity in transgenic animals. **A)** Microscope images of *hsp-6*::GFP worms obtained with brightfield (top), and fluorescence (middle) channels, and individual worm masks (bottom) segmented by the model. **B)** Median fluorescence intensity of *hsp-6*::GFP worms increases with exposure to high concentrations of zinc. **C)** Length of *hsp-6*::GFP worms decreases with exposure to high concentrations of zinc. Asterisks denote significant differences from the 0mM control: * p<1e-3; ** p<1e-4; *** p<1e-5, two-sided Welch’s t-test with Holm multiple comparison correction.

The masks also faithfully reproduce morphological differences. Zinc toxicity has prevIoUsly been shown to affect the morphology of adult worms ^33,34^. We used the predicted masks to extract length measurements of zinc exposed worms, and found that worm length decreased with high exposure to zinc (Fig 3C).

### Extending the model to additional functions

The flexible structure of the pipeline allows the user to easily adapt the model for new tasks. For example, by adding body-part specific masks to the training input the model can learn to identify that part in addition to the full worm masks. Depending on the target body part, this can be achieved with minimal additional manual labeling. To demonstrate this feature, we constructed head-specific masks by simply extracting the front 15% of each ground-truth worm mask. This required a single extra click per worm to mark the head location in the manual annotation GUI (i). The head masks were concatenated with the ground-truth labels (foreground, boundary, and seed) before training, and the model was extended with an additional output class and retrained.

With this, the inference module (iii) is able to locate the centroid of the target body part. To demonstrate the utility of this, and to showcase the generalizability of the pipeline, we applied the model to a new collection of images obtained by a third experimenter with a different acquisition protocol, including a smaller magnification. For these images we used the *hsp-4*::GFP strain, whose expression pattern is different from the *hsp-6*::GFP (Fig 4A). We then used the predicted head location (Fig 4B) to compare the fluorescence profile along the body of the worms in both strains. Figure 4C shows the difference in fluorescence intensity distribution. The *hsp-4* promoter shows two main foci of expression: a narrow peak just behind the pharyngeal bulb, and a wider peak in the posterior region, encompassing about a third of the body length. In contrast, *hsp-6* shows a single intensity peak around the anterior gut region, faithfully recreating the difference in expression patterns of the two promoters.

**Figure 4.**
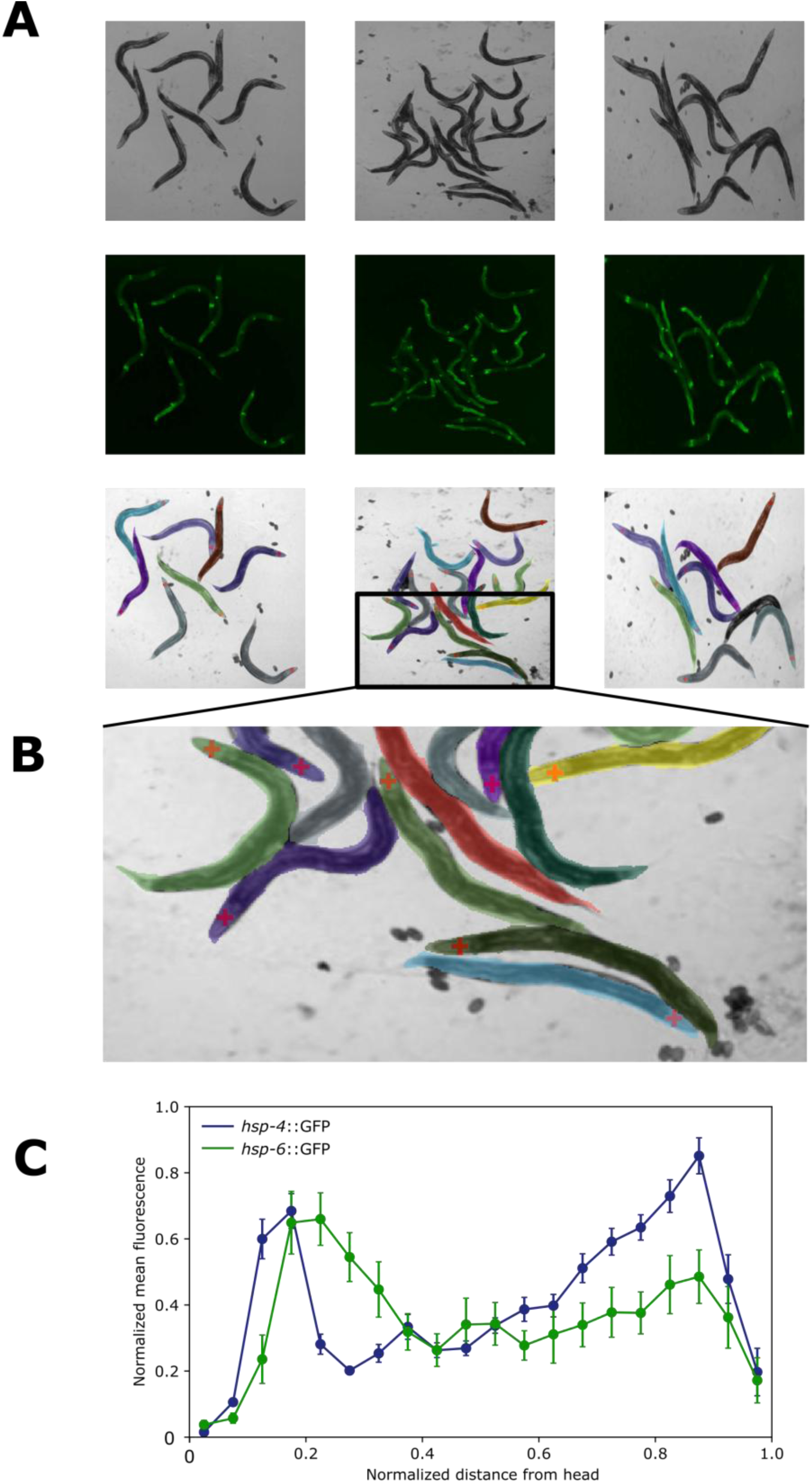
Distribution of fluorescence along the body, obtained by expanding the model to segment the head region. **A)** Microscope images of *hsp-4*::GFP worms obtained with brightfield (top), and fluorescence (middle) channels, and individual worm masks (bottom) segmented by the extended model. **B)** Zoomed in image of the worm masks showing the centroid of the segmented head region (red plus sign). **C)** Comparison of fluorescence profile of *hsp-4*::GFP vs *hsp-6*::GFP worms. Fluorescence intensity was measured in each worm in 20 bins starting from the model-identified head. Intensity was normalized within each worm before averaging.

Overall, the toolkit’s open-source and flexible architecture makes it highly generalizable, adaptable, and expandable, offering high utility to any instance segmentation task, even outside of the included pre-trained model.

## Discussion

Reliable segmentation is foundational for quantitative biological imaging, from morphological profiling and organism counting to tracking behavior and quantifying fluorescence. The field has converged on a handful of practical solution families, each reflecting a different tradeoff between speed, accuracy, and usability. Classical pipelines (thresholding, morphology, watershed) are fast and interpretable, but often brittle. Semantic segmentation models, most often U-Net, are generally more robust, but shift the burden to curating training data, compute, and maintenance, and pre-trained models may not transfer cleanly across microscopes or acquisition settings. Instance-focused methods address touching and overlap using detection-first approaches (e.g., Mask R-CNN), but these can be heavier to deploy, sensitive to domain shift, or still require manual correction in edge cases. Together, these tradeoffs highlight the need for tools that are lightweight, robust, and adaptable, while remaining approachable to non-experts.

SegBio is designed to fit in this tradeoff intersection, making instance segmentation of crowded adult *C. elegans* microscopy images practical in day-to-day experimental work and accessible to a wide user base, regardless of prior experience or hardware availability. It offers a lightweight end-to-end pipeline spanning training-label creation, model training, inference, and post-hoc correction organized into three separate but interconnected modules.

A central focus in the design is to reduce the cost of producing labels for supervised learning. Rather than requiring painstaking manual tracing of every worm, the annotation workflow (i) relies on a simple anatomical prior, the tapering profile, to extrapolate object masks from a minimal, few-click centerline. Though the resulting masks are not always pixel-perfect, this compromise preserves the benefits of supervised learning while making it realistic to generate sufficient labeled data. The flexible, heuristic-based implementation also means that the utility of the annotation tool is not confined to *C. elegans* nematodes. As long as the target objects are relatively uniform in proportions, a custom tapering profile can be provided to adapt the entire pipeline to any object or animal shape.

On the modeling side, the approach is as broad as possible. Featuring flexible model architecture, and a generalizable inference algorithm, the training module (ii) can be leveraged to accommodate similar segmentation tasks of varying complexity. However, in the spirit of accessibility, the library comes with a pre-trained plug-and-play model for brightfield adult *C. elegans* images which can be immediately utilized for a variety of experimental purposes. We demonstrate several such applications including morphological and fluorescence intensity measurements, and we are confident that the *C. elegans* community will swiftly come up with additional uses.

While the current model’s performance is generally high, some limitations remain. First, the provided pre-trained model is inherently domain-bound: performance depends on imaging modality, magnification, contrast, developmental stage, and how similar worm appearances are to the training distribution. Deviations from these parameters will require fine-tuning or retraining. Second, the current architecture does not address overlapping worms. Other tools are available that address overlaps explicitly ^35^, and perhaps future versions of SegBio will incorporate such edge cases. For this initial version, the experimenters will be required to account for worm positioning during image acquisition. Alternatively, such edge cases can be removed in the final inference step.

While the model itself is relatively simplistic, a major practical differentiator is the inference workflow. We are acutely aware that no model is perfect, and occasional errors are unavoidable. We address this issue by minimizing correction friction. Inference is performed in a completely standalone intuitive application that does not involve any complex installation procedures, has no dependencies, and does not even require an existing Python installation. The editing itself is performed on intermediate representations of the output (boundary/seed layers), followed by a repeat of post-processing, which is much faster than repeated inference or manual correction of the masks themselves.

Overall, SegBio’s main impact is shortening the path from microscopy images to clean, per-animal instance masks and measurements while keeping the workflow accessible to non-expert users. By treating segmentation as an iterative, correctable process, supported by rapid label generation, separation-aware predictions, and a packaged editor, the tool aims to make large-scale animal phenotyping more practical and scalable across experiments.

## Methods

### Strains and worm maintenance

MIR249 *risIs33* [*K03A1.5p*::3xFLAG::SV40-NLS::dCas9::SV40-NLS::VP64::HA + *unc-119*(+)] ^36^

SJ4005; *zcIs4* [*hsp-4*::GFP] V ^37^

SJ4100 (*zcIs13*[*hsp-6*::GFP]) ^38^

Worms were grown at 20°C on NGM plates seeded with *E. coli*.

### Molecular biology and RNAi

L4440-*zntA*::*cco-1* plasmid was made from L4440-*cco-1* plasmid (From the *C. elegans* RNAi library, gene name *F26E4.9* or *cox-5b* ^39^), by replacing both flanking T7 promoters with two 500bp promoters of the *E. coli zntA* gene from both sides of the *cco-1* RNAi. *E. coli* HT115 bacteria were transformed with our L4440-*zntA::cco-1* plasmid. For each experiment a fresh colony was picked and grown in LB for 8h before seeding on NGM Ampicillin plates containing increasing concentrations of ZnSO_4_. The plates were left to dry for 24 h before the experiment.

SJ4100 (*zcIs13*[*hsp-6*::GFP]) worms were bleached to obtain eggs that were placed on the HT115 seeded NGM plates. Worms were grown to the young adult stage. zntA in *E. coli* responds to Zinc in the environment ^40^. Since we used the *zntA* promoter to express RNAi directed to the worm *cco-1* gene in the HT115 bacteria, worms were fed with *cco-1* RNAi which led to a mitochondrial unfolded protein response stress and expression of GFP derived from the hsp-6 promoter.

### Image acquisition

10-15 Young adult worms were transferred onto a clean agar plate and immobilized with a drop of 10^-4^ levamisole.

All training images were acquired with a QImaging QIClick 12-bit monochrome camera attached to an Olympus MVX10 binocular, using an Olympus MV PLAPO 2XC lens and with 2.5x zoom, controlled by QCapture.

The additional test set was acquired with an Olympus IX-83 inverted microscope equipped with a Photometrics EMCCD camera and a 4x Olympus objective controlled by MicroManager.

### Manual annotation for training-label generation

To initialize the segmentation pipeline, we created a simple interactive annotation tool that enables rapid construction of training labels from raw worm images. The goal of this first stage is to convert a small amount of manual input into consistent, per-worm binary masks that can be used to train downstream deep-learning models.

### User-guided annotations

To create training masks for the model the user must provide the following:

1. Per-instance centerline (a polyline drawn along the worm’s midbody, Fig 5B)
2. Per-instance width measurement (a short cross-body line at the widest point).
3. A tapering profile of the generic object (denoted in fractions relative to max width).
4. (Optional) Per-instance head location

**Figure 5.**
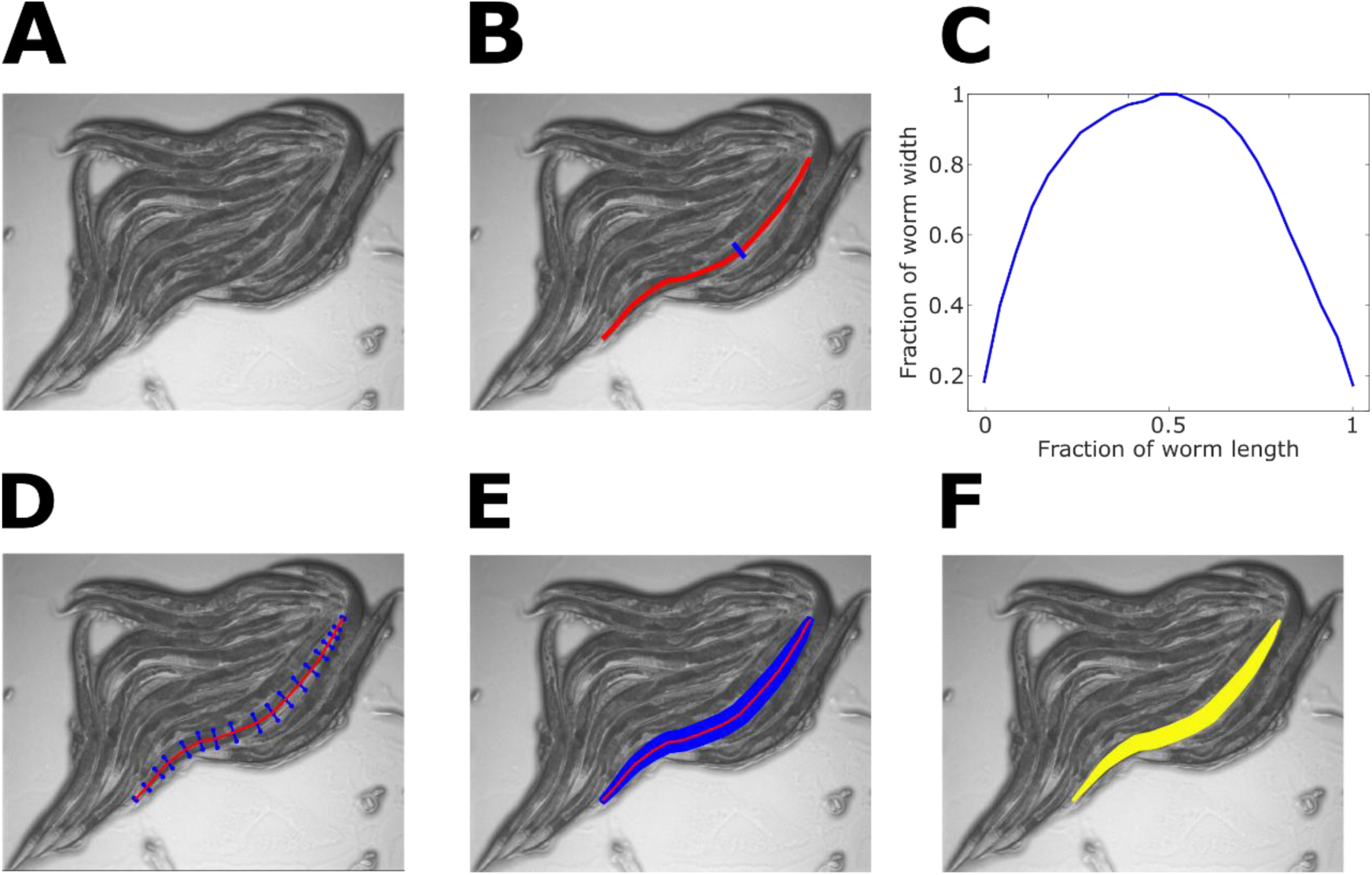
Full instance masks are extrapolated from minimal manual annotations. **A)** A brightfield image of adult C. elegans nematodes **B)** User-provided few-points centerline (red) and max width (blue) of an individual nematode **C)** Tapering profile as a fraction of max width, obtained by averaging width measurements along the bodies of 20 worms **D)** The mask is constructed by extending lines (blue) orthogonally from the centerline (red), scaled by the max width and tapering profile **E)** The centerline is interpolated to densify the mask and cover the entire worm **F)** The full mask is obtained by intersecting the orthogonal lines and filling in any remaining gaps.

A typical tapering profile of an adult *C. elegans* is provided in the code, but the pipeline can be adapted to any elongated object by a custom profile.

These sparse annotations capture the essential worm geometry while keeping annotation time low.

### Mask extrapolation from sparse annotations (centerline + width)

Each worm mask is synthesized from the user-drawn centerline by expanding outward around the midline to form a filled body region in accordance with the tapering profile (Fig X A-F). The tapering profile was obtained once by measuring the width of 20 worms at multiple points along the body, expressing width as percentage of the maximum width of each worm, and averaging (Fig 5C). The final profile is slightly asymmetric due to the different widths in the anterior and posterior regions of the worm

To form the object masks, the centerline is first densified to obtain a smooth sequence of points, and at each point the local centerline direction is estimated and a short cross-section perpendicular to the midline is constructed (Fig 5D,E). The length of these cross-sections is scaled by the user’s width measurement and modulated by the tapering profile, so the mask gradually narrows toward the head and tail. The union of all cross-sections is then converted into a single contiguous binary region and lightly regularized (gap-closing) to yield the final per-worm pixel mask (Fig 5F). This produces anatomically plausible masks from a single width measurement while still allowing variation across worms via the user-provided width.

For each image, the software saves the user annotations (centerline and width) together with the derived outputs (per-worm binary masks and basic geometry summaries). These outputs serve as the ground-truth labels for training the segmentation model in subsequent stages.

The annotation tool was originally written as a MATLAB (MathWorks^©^ Inc.) app, and later rewritten and expanded in Python. Both versions are available at https://github.com/zaslab/SegBio.

## Model training pipeline

### Training target generation from instance masks

To train the network with supervision that supports both object presence and instance separation, the manually created per-worm instance mask (integer label image; background = 0, worm IDs = 1. . *N*) is converted into three complementary target maps: foreground, boundary, and seed (Fig X D-H) to be used as ground truth during model training.

### Foreground target

A binary foreground map is produced directly from the instance labels providing dense supervision for “worm vs background” classification.

### Boundary target

To encourage separation of touching or nearby worms, a thin boundary band is computed along the outer rim of each labeled instance (and between neighbors). This rim is optionally dilated to a user-defined pixel width (boundary_width) to form a thicker supervision band. The boundary target is restricted to foreground pixels to avoid labeling background edges unrelated to worms.

### Seed target

In addition to boundaries, a seed target is generated to provide one compact “core” signal per worm instance, intended to anchor downstream instance reconstruction.

The default seed method derives a per-instance skeleton, prunes a small fraction from each end to avoid ambiguous head/tail tips, then slightly thickens the skeleton to form a learnable region. Seed pixels overlapping the boundary band are removed, and small spurIoUs components are filtered by a minimum-area criterion to preserve only robust seed blobs.

As an alternative, seeds can be generated from the distance transform of the foreground mask, producing a soft interior-confidence map that peaks near the center of worms; boundary-adjacent pixels are suppressed so the seed emphasizes the worm interior.

The output is a dictionary containing the three aligned targets:

- fg: foreground worm mask
- boundary: boundary band for separation
- seed: per-instance interior anchors (skeleton-based by default)

### Neural network architecture

The model is a parameterized U-Net–style fully convolutional network that predicts the training targets from input microscopy images. The model is designed to be lightweight, configurable (depth, channel width, output classes), and stable under small batch sizes to accommodate less powerful hardware.

### Overall structure

The network follows the standard encoder–bottleneck–decoder U-Net topology:

- Encoder (contracting path): repeated blocks of two 3 × 3 convolutions with normalization and ReLU, followed by 2 × 2 max-pooling for downsampling. Feature channel count doubles with depth.
- Bottleneck: a double-convolution block at the deepest level with 2x the channels of the deepest encoder stage.
- Decoder (expanding path): transposed-convolution upsampling, concatenation with the corresponding encoder feature map via skip connections, then a double-convolution block to fuse features.

This symmetric design preserves fine spatial detail (via skips) while still learning high-level context (via downsampling).

Each convolutional block is a DoubleConv module implementing:

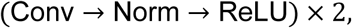

with 3 × 3 convolutions (padding to preserve spatial size) and Group Normalization. The number of groups is automatically chosen as the largest value ≤ 8 that divides the channel count, improving robustness when batch size is small.

A final 1 × 1 convolution maps decoder features to the n_classes output channels. During upsampling, feature maps are padded when needed so that the upsampled tensor matches the skip-connection tensor size, ensuring correct concatenation even for odd image dimensions.

### Configurable capacity

The training code is designed to allow for quick and easy adaptation of the model to new datasets. Users can provide their own annotated images, or annotate images themselves using the annotation module (i), and retrain the model to better fit the new data. Model capacity can be adjusted to the complexity of the new task by controlling the following parameters:

- depth: number of encoder/decoder levels (≥2),
- base_filters: number of channels at the first level, with doubling at each deeper level,
- n_channels: input channels (e.g., brightfield only or multi-channel),
- n_classes: output channels, enabling additional segmentation targets such as head regions.

### Dataset format and preprocessing

Training samples are stored as per-image folders containing a raw input image (in.mat) and a corresponding manual annotation mask (out.mat). During loading, inputs and masks are resized to a fixed 512 × 512 resolution (image interpolation preserves intensity, while masks are interpolated with nearest-neighbor to preserve binary labels), and image intensities are normalized to [0,1].

### Data augmentation (supported transforms)

Training uses an augmentation module that applies the same geometric transform to the input image and all target channels and applies photometric transforms to the image only.

Geometric (image + targets, shared per sample):

- Random zoom-out / scale-down with random placement on a canvas (acts like zoom + random padding/positioning).
- Horizontal flip and vertical flip (each with 0.5 probability).
- Random affine transform consisting of rotation (±rot_deg) and a small-scale jitter (≈0.95–1.05).

Photometric (image only):

- Brightness jitter (multiplicative intensity scaling).
- Contrast jitter (contrast change around the per-channel mean).
- Gaussian blur applied with probability blur_p.

Multi-augmentation per batch: the training loop can generate multiple independent augmentations of each minibatch and accumulate gradients across them prior to the optimizer step.

### Loss function

Training optimizes a multi-task objective combining Binary Cross Entropy (BCE) and Dice, with class weights of [1,6,2] for foreground, boundary, and seed, respectively. The final loss is the weighted sum *BCE* + 0.5 × *Dice*.

### Optimization, scheduling, and validation

Models are trained using the AdamW optimizer with weight decay and a step learning-rate schedule. Training optionally uses mixed precision (AMP) via gradient scaling, and can optionally use torch.compile for accelerated execution. A held-out validation split is created once at the beginning, and training logs loss/Dice metrics to a CSV file while saving periodic checkpoints.

### Post processing

At inference time, the network produces three probability maps (foreground, boundary, seed). These are binarized by thresholds and then cleaned using connected-component filtering to suppress small fragments and retain only substantial regions. Boundary maps are additionally thinned to sharpen separation cues.

Instance labels are obtained with a marker-controlled watershed over a distance-based (or probability-based) surface, constrained to the cleaned foreground region and discouraged from merging across the predicted boundaries. This converts the three maps into a final integer label image where each worm is a distinct instance.

After instance generation, labels can be filtered using skeleton-derived morphology: per-instance skeleton length, width estimate (via medial axis distance), and length/width ratio are computed and used to remove implausible detections. An optional rule removes instances touching the image border, avoiding partial objects. Remaining instances are relabeled into a compact 1. . *N* index set.

### Interactive inference and manual correction GUI

Inference is performed with a standalone graphical application that couples the trained network with an interactive editing workflow. The purpose of this stage is to (i) make segmentation usable for non-expert users, and (ii) ensure high-quality final labels in difficult frames by allowing fast, local corrections to either the model output or the final inference masks.

### Model execution and initial segmentation

After opening an image, the application runs the network once to generate three outputs: a foreground probability map, and two auxiliary maps used for instance separation (boundary and seed). These are thresholded to initialize editable “boundary” and “seed” layers, and an initial set of instance labels is generated from the three-map representation.

### Human-in-the-loop editing

The GUI exposes the network-derived boundary and seed maps as paintable layers. This design makes edits intuitive:

- Fix merges by painting boundary lines between worms to force separation.
- Fix splits or missing worms by removing extra boundaries or adding seed regions to missed worms
- Delete false positives with a dedicated button by clicking an unwanted instance label
- Manually paint instance masks for ultimate control and precision

Edits are lightweight and local, and do not require re-running the neural network.

After edits, the user triggers “Re-segment,” which recomputes the instance labels using the original predicted foreground together with the edited boundary and seed layers. This allows rapid iteration: edit → re-segment → inspect, until the instance labels are satisfactory. A small threshold control is also provided to adjust sensitivity when foreground is under- or over-inclusive. The GUI exports the final instance label image along with the edited boundary/seed layers and basic per-instance size summaries.

A detailed user guide is provided in https://github.com/zaslab/SegBio.

### Training Hardware

The entire pipeline was designed to be as lightweight and accessible as possible, without relying on extensive programming knowledge, or high-end hardware for either training or inference.

The model was trained on the following hardware:

- Lenovo Legion 5 laptop
- Intel(R) Core(TM) i7-14650HX (2.20 GHz) processor
- 32GB RAM
- Nvidia GeForce RTX 4060 laptop GPU, 8GB VRAM

With a depth of 4, filter count of 32, a batch size of 6, and 5 augmentations per image training engaged most of the available VRAM. Training time was ∼7 seconds per batch. With a total of 175 training images, a full training run of 100 epochs can be completed in under 5 hours.

### Funding

This work was supported by the Israeli Science Foundation (1939/23), the Jérôme Lejeune Foundation, and the MDBR grant program of the Orphan Disease Center. AZ is the Greenfield Chair in Neurobiology.

### Data availability

All data, analysis scripts, and the entire segBio package is available in https://github.com/zaslab/SegBio.

## References

1. Ronneberger, O., Fischer, P. & Brox, T. U-Net: Convolutional Networks for Biomedical Image Segmentation. in Medical Image Computing and Computer-Assisted Intervention – MICCAI 2015 (eds Navab, N., Hornegger, J., Wells, W. M. & Frangi, A. F.) vol. 9351 234–241 (Springer International Publishing, Cham, 2015).

2. Falk, T. et al. U-Net: deep learning for cell counting, detection, and morphometry. Nat. Methods 16, 67–70 (2019).

3. Galimov, E. & Yakimovich, A. A tandem segmentation-classification approach for the localization of morphological predictors of C. elegans lifespan and motility. Aging 14, (2022).

4. He, K., Gkioxari, G., Dollár, P. & Girshick, R. Mask R-CNN. Preprint at 10.48550/arXiv.1703.06870 (2018).

5. Dong, B. & Chen, W. A high precision method of segmenting complex postures in Caenorhabditis elegans and deep phenotyping to analyze lifespan. Sci. Rep. 15, 8870 (2025).

6. Liu, X. et al. Automated C. elegans behavior analysis via deep learning-based detection and tracking. PLOS Comput. Biol. 21, e1013707 (2025).

7. Schmidt, U., Weigert, M., Broaddus, C. & Myers, G. Cell Detection with Star-Convex Polygons. in Medical Image Computing and Computer Assisted Intervention – MICCAI 2018 (eds Frangi, A. F., Schnabel, J. A., Davatzikos, C., Alberola-López, C. & Fichtinger, G.) vol. 11071 265–273 (Springer International Publishing, Cham, 2018).

8. Mais, L., Hirsch, P. & Kainmueller, D. PatchPerPix for Instance Segmentation. in Computer Vision – ECCV 2020 (eds Vedaldi, A., Bischof, H., Brox, T. & Frahm, J.-M.) vol. 12370 288–304 (Springer International Publishing, Cham, 2020).

9. Deserno, M. & Bozek, K. WormSwin: Instance segmentation of C. elegans using vision transformer. Sci. Rep. 13, 11021 (2023).

10. Stringer, C., Wang, T., Michaelos, M. & Pachitariu, M. Cellpose: a generalist algorithm for cellular segmentation. Nat. Methods 18, 100–106 (2021).

11. Cutler, K. J. et al. Omnipose: a high-precision morphology-independent solution for bacterial cell segmentation. Nat. Methods 19, 1438–1448 (2022).

12. Vincent, L. & Soille, P. Watersheds in digital spaces: an efficient algorithm based on immersion simulations. IEEE Trans. Pattern Anal. Mach. Intell. 13, 583–598 (1991).

13. Naylor, P., Laé, M., Reyal, F. & Walter, T. Segmentation of Nuclei in Histopathology Images by Deep Regression of the Distance Map. IEEE Trans. Med. Imaging 38, 448–459 (2019).

14. Xie, L., Qi, J., Pan, L. & Wali, S. Integrating deep convolutional neural networks with marker-controlled watershed for overlapping nuclei segmentation in histopathology images. Neurocomputing 376, 166–179 (2020).

15. Shah, P., Bao, Z. & Zaidel-Bar, R. Visualizing and quantifying molecular and cellular processes in *Caenorhabditis elegans* using light microscopy. Genetics 221, iyac068 (2022).

16. Wählby, C. et al. An image analysis toolbox for high-throughput C. elegans assays. Nat. Methods 9, 714–716 (2012).

17. Weheliye, W. H. et al. A neural network model enables worm tracking in challenging conditions and increases signal-to-noise ratio in phenotypic screens. PLOS Comput. Biol. 21, e1013345 (2025).

18. Sydney Brenner. The Genetics of Caenorhabditis elegans. Genetics 71–94 (1974) doi:10.1093/genetics/77.1.71.

19. Barlow, I. L. et al. Megapixel camera arrays enable high-resolution animal tracking in multiwell plates. *Commun*. Biol. 5, 253 (2022).

20. Ambros, V. R., et al. From nematode to Nobel: How community-shared resources fueled the rise of *Caenorhabditis elegans* as a research organism. Proc. Natl. Acad. Sci. 122, e2522808122 (2025).

21. Min, H., Park, G. & Lee, S.-J. V. Brief guide to Caenorhabditis elegans imaging and quantification. Mol. Cells 48, 100249 (2025).

22. Ji, H., Dian, Chen, & Fang-Yen, Christopher. Automated multimodal imaging of Caenorhabditis elegans behavior in multi-well plates. Genetics 228, (2024).

23. Baek, J.-H., Cosman, P., Feng, Z., Silver, J. & Schafer, W. R. Using machine vision to analyze and classify Caenorhabditis elegans behavioral phenotypes quantitatively. J. Neurosci. Methods 118, 9–21 (2002).

24. Feng, Z., Cronin, C. J., Wittig, J. H., Sternberg, P. W. & Schafer, W. R. An imaging system for standardized quantitative analysis of C. elegans behavior. BMC Bioinformatics 5, 115 (2004).

25. Stephens, G. J., Johnson-Kerner, B., Bialek, W. & Ryu, W. S. Dimensionality and Dynamics in the Behavior of C. elegans. PLOS Comput. Biol. 4, e1000028 (2008).

26. Itskovits, E., Levine, A., Cohen, E. & Zaslaver, A. A multi-animal tracker for studying complex behaviors. BMC Biol. 15, 29 (2017).

27. Ljosa, V., Sokolnicki, K. L. & Carpenter, A. E. Annotated high-throughput microscopy image sets for validation. Nat. Methods 9, 637–637 (2012).

28. Escobar-Benavides, S., García-Garví, A., Layana-Castro, P. E. & Sánchez-Salmerón, A.-J. Towards generalization for Caenorhabditis elegans detection. Comput. Struct. Biotechnol. J. 21, 4914–4922 (2023).

29. Venkatachalam, V. et al. Pan-neuronal imaging in roaming *Caenorhabditis elegans*. Proc. Natl. Acad. Sci. 113, (2016).

30. Yu, X. et al. Fast deep neural correspondence for tracking and identifying neurons in C. elegans using semi-synthetic training. eLife 10, e66410 (2021).

31. Park, C. F. et al. Automated neuron tracking inside moving and deforming C. elegans using deep learning and targeted augmentation. Nat. Methods 21, 142–149 (2024).

32. Sprague, D. Y. et al. Unifying community whole-brain imaging datasets enables robust neuron identification and reveals determinants of neuron position in C. elegans. *Cell Rep*. Methods 5, 100964 (2025).

33. Khare, P. et al. Size dependent toxicity of zinc oxide nano-particles in soil nematode *Caenorhabditis elegans*. Nanotoxicology 9, 423–432 (2015).

34. Moyson, S., Town, R. M., Vissenberg, K. & Blust, R. The effect of metal mixture composition on toxicity to C. elegans at individual and population levels. PLOS ONE 14, e0218929 (2019).

35. Castro, P. E. L. et al. SegElegans: Instance segmentation using dual convolutional recurrent neural network decoder in Caenorhabditis elegans microscopic images. Comput. Biol. Med. 190, 110012 (2025).

36. Fischer, F. et al. Ingestion of single guide RNAs induces gene overexpression and extends lifespan in Caenorhabditis elegans via CRISPR activation. J. Biol. Chem. 298, 102085 (2022).

37. Kapulkin, V., Hiester, B. G. & Link, C. D. Compensatory regulation among ER chaperones in *C. elegans*. FEBS Lett. 579, 3063–3068 (2005).

38. Yoneda, T. et al. Compartment-specific perturbation of protein handling activates genes encoding mitochondrial chaperones. J. Cell Sci. 117, 4055–4066 (2004).

39. Kamath, R. S. et al. Systematic functional analysis of the Caenorhabditis elegans genome using RNAi. Nature 421, 231–237 (2003).

40. Durieux, J., Wolff, S. & Dillin, A. The Cell-Non-Autonomous Nature of Electron Transport Chain-Mediated Longevity. Cell 144, 79–91 (2011).

